# Cryptic Native American ancestry recapitulates population-specific migration and settlement of the continental United States

**DOI:** 10.1101/333609

**Authors:** I. King Jordan, Lavanya Rishishwar, Andrew B. Conley

## Abstract

European and African descendants settled the continental US during the 17^th^-19^th^ centuries, coming into contact with established Native American populations. The resulting admixture among these groups yielded a significant reservoir of cryptic Native American ancestry in the modern US population. We analyzed the patterns of Native American admixture seen for the three largest genetic ancestry groups in the US population: African American, European American, and Hispanic/Latino. The three groups show distinct Native American ancestry profiles, which are indicative of their historical patterns of migration and settlement across the country. Native American ancestry in the modern African American population does not coincide with local geography, instead forming a monophyletic group with origins in the southeastern US, consistent with the Great Migration of the early 20^th^ century. European Americans show Native American ancestry that tracks their geographic origins across the US, indicative of ongoing contact during westward expansion, and Native American ancestry can resolve Hispanic/Latino individuals into distinct local groups formed by more recent migration from Mexico and Puerto Rico. We found an anomalous pattern of Native American ancestry from the US southwest, which most likely corresponds to the *Nuevomexicano* descendants of early Spanish settlers to the region. We addressed a number of controversies surrounding this population, including the extent of Sephardic Jewish ancestry. *Nuevomexicanos* are less admixed than nearby Mexican-American individuals, with more European and less Native American and African ancestry, and while they do show demonstrable Sephardic Jewish ancestry, the fraction is no greater than seen for other Hispanic/Latino populations.

## Introduction

Native Americans inhabited the area that now makes up the continental US for thousands of years prior to the arrival of the first European settlers. The ancestors of modern Native Americans are thought have arrived in the Americas from Asia, by way of the Bering Strait, in several successive waves of migration^1^. The current model, based on archaeology and comparative genomic studies, holds that the earliest ancestors of Native Americans arrived in the Americas ∼23,000 years ago^2^. The earliest evidence for Native Americans in the continental US dates to ∼14,000 years ago^3^. The much later arrival of Europeans in the Americas, followed shortly thereafter by Africans who were brought by force via the trans-Atlantic slave trade, had a drastic effect on the demographic makeup of the region. Native American population numbers declined rapidly in the face of continuous immigration, settlement, and conflict, and as a result the modern US population is made up mainly of descendants of European and African immigrants.

Europeans arrived in the Americas more than 20,000 years after the first Native Americans. The first European settlers to reach the continental US were Spaniards led by the conquistador Ponce de León, who claimed Florida for the Spanish crown in 1513^4^. British settlers arrived more than 70 years later, initially establishing the ill-fated colony of Roanoke in 1585 and later the permanent settlement of Jamestown in 1607^5^. An estimated 400,000 British had migrated to the US by the end of the 17^th^ century. The first Africans were brought to Jamestown in 1619 by Dutch pirates who traded them to the British settlers as indentured servants^6^. The social status of Africans in the US changed quickly, with slavery first legally sanctioned by 1640. The trans-Atlantic slave trade would eventually bring ∼400,000 enslaved Africans to the continental US^7^.

The arrival of Europeans and Africans in the Americas, and the conflict that followed, would prove to be catastrophic for the indigenous population. It has been estimated that 10-100 million Native Americans may have died in the first 150 years after Columbus’ arrival in the New World, amounting to a 95% reduction in the population^8^. This massive Native American population decline is mainly attributed to the introduction of European and African endemic infectious diseases – *e.g.* malaria, measles, and smallpox – for which the indigenous population had little or no immune defense.

The story of conflict between Native Americans and European and African settlers, along with the devastating consequences for the indigenous population, is by now well-known. However, there is another, perhaps less appreciated, aspect of the encounter between these population groups that has also had profound consequences for the genetic demography of the Americas. Here, we are referring to the process of genetic admixture, whereby individuals from previously isolated population groups reproduce, resulting in the combination of ancestry-specific haplotypes within individual genomes. Admixture has been a fundamental feature of human evolution and migration^9^. Whenever previously isolated human populations meet, no matter what the circumstances, they mix and give rise to individuals with a mosaic of different genetic ancestries.

As European and African descendants settled the continental US, they inevitably came into contact with established Native American populations resulting in admixture and the introgression of Native American genomic sequence into the expanding US population. Accordingly, the genomes of European and African descendants in the US are expected to contain some fraction of Native American ancestry. In other words, a significant reservoir of Native American ancestry currently exists outside of recognized indigenous communities. We refer to this ancestry component as ‘cryptic’ Native American ancestry given the fact that its low levels may often lead it to go unrecognized. In this study, we ask how the historical processes of migration and settlement affected the distribution of cryptic Native American ancestry across the continental US. We address this question for the three largest genetic ancestry groups in the modern US population: African American, European American, and Hispanic/Latino.

## Material and Methods

### Genotype datasets

Whole genome genotype data of US individuals from the Health and Retirement Study (HRS) dataset (*n*=15,620) were merged with whole genome sequence variant data from the 1000 Genomes Project (1KGP)^10;11^ (*n*=1,718) and whole genome genotype data from the Human Genome Diversity Project (HGDP)^12–14^ (*n*=230) (Table S1). HRS genotype data were accessed via the NCBI dbGaP database and the study was conducted with Institutional Review Board approval from the Georgia Institute of Technology (protocol number H17029). Individual HRS genotypes are provided along with geographical origin data for sample donors from the nine census regions in the continental US. A collection of Native American genotypes from 21 populations across the Americas was taken from a comprehensive study on Native American population history^2^ (*n*=314). These Native American genotype data were accessed according to the terms of a data use agreement from the Universidad de Antioquia. Whole genome genotype data from 5 populations of Sephardic Jewish individuals (*n*=40) were also included as reference populations^15^. The genotypes from HRS individuals were merged with the comparative genomic data sources using PLINK version 1.9^16^, keeping only those sites common to all datasets and correcting SNP strand orientations for consistency as needed. The final merged dataset includes 228,190 SNPs across 17,882 individuals. The merged genotype dataset was phased using ShapeIT version 2.r837^17^. SNPs that interfered with the ShapeIT phasing process were excluded from subsequent analyses. ShapeIT was run without reference haplotypes, and all individuals were phased at the same time. Individual chromosomes were phased separately, and the X chromosome was phased with the additional ‘-X’ flag.

### Local ancestry inference

The RFMix algorithm^18^ is able to accurately characterize the local ancestry of admixed individuals but is prohibitively slow when run on a dataset of the size used here. To reduce the runtime, we modified RFMix version 1.5.4 so that the expectation-maximization (EM) procedure samples from, and creates a forest for, the entire set of individuals rather than each individual. This modified RFMix was run in the PopPhased mode with a minimum node size of five, using 12 generations and the “--use-reference-panels-in-EM” for two rounds of EM, generating local ancestry inference for both the reference and admixed populations. Continental African, European, and Native American populations were used as reference populations. Contiguous regions of ancestral assignment, “ancestry tracts,” were created where RFMix ancestral certainty was at least 95%. Genome-wide ancestry estimates from the modified RFMix algorithm closely correlate with those from ADMIXTURE (Figure S1).

The extent of Sephardic Jewish (*Converso*) ancestry in individuals from the Hispanic/Latino group in HRS (as defined in the genome-wide ancestry section below), and Latin American populations from 1KGP, was inferred via ancestry-specific haplotype comparisons with Sephardic Jewish reference populations using the program ChromoPainter2^9^ (kindly provided by Garrett Hellenthal). First, African and Native American haplotypes were masked from the RFMix output. Then, the remaining European haplotypes were compared against genomes from the European reference populations together with the Sephardic Jewish populations. The extent of Jewish ancestry for any individual genome is defined as the ‘copying fraction’ from the Sephardic Jewish populations, where the copying fraction is taken as the fraction of sites with best matches to the Sephardic Jewish reference genomes. It should be noted that this procedure results in a relative fraction of Sephardic Jewish ancestry for all individuals under consideration, which is directly comparable among individuals but likely to be an overestimate of the total ancestry derived from a single source population.

### Genome-wide ancestry inference

ADMIXTURE^19^ version 1.3.0 was used with *K*=4 to infer continental ancestry fractions for individuals in the dataset via comparison with reference populations from Africa, Europe, the Americas, and East Asia. Sub-continental ancestry was inferred independently for each of the three major continental ancestry components – African, European, and Native American – using an ancestry-specific masking procedure that we developed as previously described^20^. This procedure relies on the local continental ancestry assignments, along with the re-phased genotypes, generated by RFMix as described above. Sub-continental ancestry was characterized by first masking out two of the three continental ancestries (African, European, and/or Native American) at a time and then analyzing the genomic regions (haplotypes) corresponding to the remaining continental ancestry. For sub-continental ancestry analysis of any given continental ancestry component, only those individuals with at least 1.5% genome-wide ancestry for that same continental group were used.

We developed a novel machine learning based approach to distinguish Spanish from other (primarily Western) European descendants in the HRS dataset via analysis of European-specific haplotypes. First, ADMIXTURE was run with *K*=5 on the RFMix characterized European haplotypes for the HRS individuals to stratify sub-continental European ancestries based on comparison with Northern (Finnish and Russian), Western (French and British), Spanish, and Southern (Italian and Sardinian) European reference populations from the 1KGP and HGDP datasets. A Support Vector Machine (SVM) classifier^21^ was then trained using the resulting ADMIXTURE ancestry vectors for the European reference populations from the four sub-continental groups: Northern, Western, Spanish, and Southern. The European-specific ADMIXTURE ancestry vectors for the HRS individuals were then classified into one of the four European sub-continental groups defined by the SVM classifier. A confidence threshold of 0.8 was used for sub-continental group assignments. For the purpose of analysis here, we consider two major groups of European descendants in the HRS data set: Spanish descendants and all others. We refer to Spanish descendants as Hispanic/Latino (HL). Non-Spanish HRS individuals with <5% African ancestry are defined as European American, whereas non-Spanish HRS individuals with at least 20% African ancestry were defined as African American (see Supplementary Methods for additional details).

### Sex-biased ancestry inference

Sex-biased ancestry contributions were inferred by comparing the RFMix characterized fractions of each continental ancestry component on the X chromosomes versus the autosomes as previously described^22;23^. For each individual genome, and each ancestry component, the normalized difference between the X chromosome ancestry fraction and the autosomal ancestry fraction (*ΔAdmix*) is defined as:

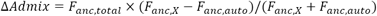

where *F*_*anc,total*_, *F*_*anc,X*_, and *F*_*anc,auto*_ are the genome-wide, X chromosome, and autosome ancestry fractions, respectively.

### Phylogenetic inference

We used the RFMix defined Native American haplotypes for individuals from the HRS and reference populations to infer the phylogenetic relationships between populations. For each pair of populations with at least 25 Native American sites covered by 5 individuals, we computed the weighted F_ST_ between all pairs of populations using PLINK. The resulting F_ST_ distance matrix was used to create a neighbor-joining tree^24^ with the program MEGA6^25^.

## Results

### Genetic ancestry groups in the US

The first aim of our study was to objectively define the major genetic ancestry groups for the continental US based on observable patterns of ancestry and admixture seen for the 15,620 HRS genotypes analyzed here. Having defined the US genetic ancestry groups, we then considered the distribution of Native American ancestry within and between ancestry groups and among geographic regions. We provide a detailed description, along with supporting results (Supplementary Figures S2-S5), of how we defined the three main US ancestry groups – African American, European American, and Hispanic/Latino – in the Supplementary Material.

The distribution of HRS individuals among the three major US genetic ancestry groups is shown in Figure 1. Visual inspection of the continental ancestry fractions seen for members of the three groups supports our objective approach to genetic ancestry-based classification (Figure 1A). For example, the majority of Hispanic/Latino individuals show substantial levels of Native American ancestry compared to individuals from the European American ancestry group (Figure 1A); the median Native American ancestry for the Hispanic/Latino group is 38% compared to 0.1% for the European American group (Figure 1B). In addition, individuals from the Hispanic/Latino group cluster tightly with the Mexican reference population from the 1KGP, along the second axis between the European and Native American populations in the principal components analysis (PCA) plot of the pairwise genome distances (Figure 1C). It is important to note that we did not use Native American ancestry for the purposes of classification. Rather, European ancestry alone was sufficient to recapitulate known levels of Native American ancestry for Hispanic/Latino individuals.

**Figure 1.**
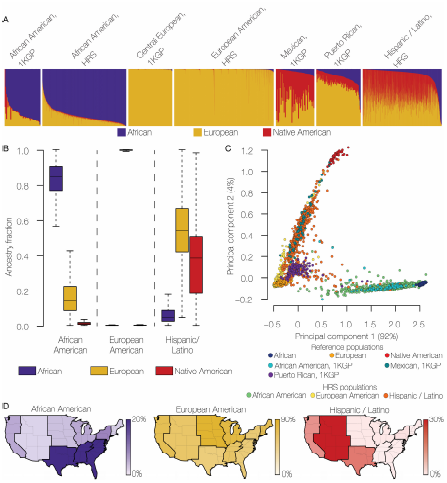
Genetic ancestry groups in the modern US population. (A) ADMIXTURE plot (*K*=3) showing the African (blue), European (gold), and Native American (red) ancestry components for individuals from different US population groups. Data are from the 1000 Genomes Project (1KGP) and the Health and Retirement Study (HRS). (B) Distributions of African, European, and Native American ancestry fractions for the three main US genetic ancestry groups defined here: African American, European American, and Hispanic/Latino. (C) Principal components analysis (PCA) plot showing the relationships among individuals from reference populations and individuals from the HRS dataset corresponding to the three US genetic ancestry groups. (D) Percentages of individuals from each of the three US genetic ancestry groups are shown for the nine census regions in continental US.

Individuals from the African American ancestry group show medians of 85% African ancestry, 14% European ancestry, and 1% Native American ancestry (Figure 1B). Most of these individuals group along the first PCA axis separating the African and European reference populations. In contrast to the admixed Hispanic/Latino and African American ancestry groups, individuals from the European American ancestry group show extremely low levels of admixture with non-European populations, with a median value of 99.8% European ancestry. Given their relatively low numbers (see Supplementary Figure S2), as well as their relatively late historical arrival in the continental US, we did not consider Asian Americans further in this study.

Individuals assigned to the three main genetic ancestry groups show distinct geographic distributions across the continental US, which are largely consistent with demographic data for the country. African ancestry is highest in the three southern census regions, European ancestry is highest in the two north central regions, and Hispanic/Latino ancestry is highest in the Mountain census region, which includes Arizona and New Mexico (Figure 1D).

### Sex-biased admixture in US genetic ancestry groups

We compared the patterns and extent of sex-biased admixture among the three US genetic ancestry groups by comparing the continental ancestry fractions – African, European, and Native American – seen for the X chromosomes versus the autosomes. For any given ancestry component, a relative excess of X chromosome ancestry is indicative of female-biased admixture, whereas an excess of autosomal ancestry reflects male-biased admixture^26^. This was only done for admixed individuals that had two or more continental ancestry fractions at >1.5% of the overall ancestry. Almost all individuals from the African American and Hispanic/Latino groups met this criterion, but only a small minority of European American individuals with Native American admixture did. African American and Hispanic/Latino ancestry groups showed marked patterns of sex-biased admixture, whereas the European Americans did not show any appreciable evidence of sex-biased admixture (Figure 2). The strongest pattern of sex-biased admixture was seen for Hispanic/Latino individuals, with female-biased Native American admixture and male-biased European admixture. African Americans show female-biased African ancestry and male-biased European ancestry.

**Figure 2.**
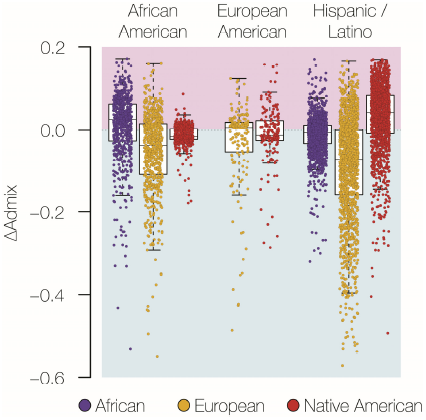
Sex-biased admixture in US genetic ancestry groups. Normalized differences between X chromosome ancestry fractions and autosomal ancestry fractions (*ΔAdmix*) are shown on the y-axis. *ΔAdmix* values are shown for each ancestry component – African (blue), European (gold), and Native American (red) – in each individual genome. *ΔAdmix* values above zero (pink) indicate female-biased admixture, and values below zero (blue) indicate male-biased admixture.

### Native American ancestry distribution across the US

For each US genetic ancestry group, we considered three distinct characteristics of Native American ancestry across the continental US: (1) the relative levels of Native American ancestry genome-wide, (2) the patterns of Native American allele frequencies, and (3) the phylogenetic relationships among US populations based on their Native American ancestry.

As we showed previously, overall Native American ancestry is highest for the Hispanic/Latino group (median 38%), followed by the African American (1%) and European American groups (0.1%) (Figure 1B). Among all three ancestry groups, the highest levels of Native American ancestry are seen for the West-South-Central (WSC; including Texas), Pacific (PAC; including California), and Mountain (MNT; including Arizona and New Mexico) census regions (Figure 3). Native American ancestry levels show the highest variability among regions for the Hispanic/Latino group (coefficient of variation [c.v.]=1.08), followed by the European American (c.v.=0.65) and then African American (c.v.=0.60) groups.

**Figure 3.**
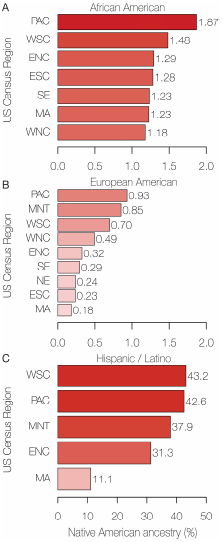
Native American ancestry percentages in the modern US population. The average percentages of Native American ancestry are shown for the three US genetic ancestry groups across the nine geographic census regions (see Supplementary Figure S3). Data for census regions with less than five individuals for any ancestry group are considered unreliable and are not shown.

We measured the patterns of Native American allele frequencies across the continental US using ADMIXTURE analysis of Native American haplotypes for individuals from the three ancestry groups. Visualization of the ancestry vectors produced by ADMIXTURE shows that the African American and European American groups have patterns that are similar to each other (Figure 4A; top panel) and distinct from the patterns seen for the Hispanic/Latino group (Figure 4B; top panel). Furthermore, the African American and European American groups show ancestry patterns that are intermediate to the Canadian (Chipewan, Algonquin, Cree, and Ojibwa) and Northern Mexican (Pima and Tepehuano) Native American reference populations, whereas the Hispanic/Latino group shows Native American ancestry patterns that are more similar to either the admixed Mexican and Native American reference populations or the admixed Puerto Rican population. There is substantially more regional variation in Native American ancestry seen for the Hispanic/Latino group, with characteristically Mexican patterns seen in the Pacific (PAC) and West South-Central (WSC) regions and a strongly Puerto Rican pattern in the Mid-Atlantic (MA) region. The Mountain region (MNT) shows a distinct and highly variable pattern of Native American ancestry, which we explore in more detail in the following section.

**Figure 4.**
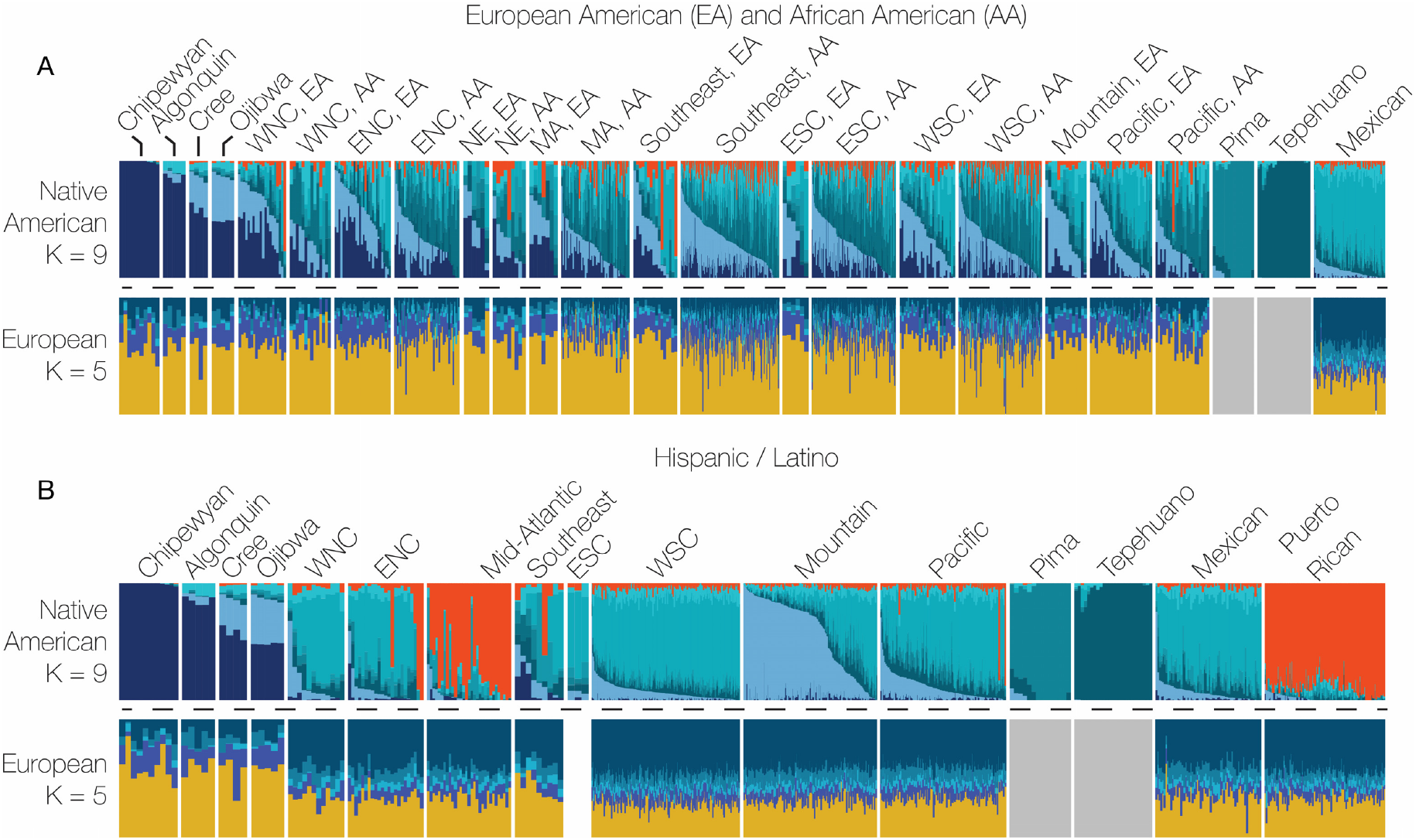
Native American and European ancestry profiles for US ancestry groups. Native American (*K*=9) and European (*K*=5) ancestry-specific ADMXITURE plots are shown for the European American (EA) and African American (AA) groups combined (A) and for the Hispanic/Latino group (B). The individual panels shown correspond to Native American reference populations (Chipewyan, Algonquin, Cree, Ojibwa, Pima, Tepehuano), 1000 Genomes Project reference populations (Mexican and Puerto Rican) and the HRS data from the different US census regions (see Supplementary Figure S3).

The phylogenetic relationships among genetic ancestry groups across the US were inferred by calculating the fixation index (F_ST_) between pairs of populations based on Native American haplotypes (Figure 5). The Canadian and Amazonian Native American reference populations occupy the most distant clades on the phylogeny with the admixed Mexican and Mexican Native American reference populations adjacent to the Amazonian group. African Americans from all of the census regions form a single monophyletic clade, with the European Americans from the Southeast region (SE) as the closest sister taxon. European Americans from the West North-Central (WNC) and East North-Central (ENC) regions group most closely with the Canadian Native American reference populations, and the European Americans from this region form a distinct group adjacent to a Mexican group of populations. Members of the Hispanic/Latino ancestry group from most of the census regions group closely with Mexican populations, with the exception of the Mid-Atlantic region (MA) which groups most closely with the admixed Puerto Rican and Amazonian reference populations.

**Figure 5.**
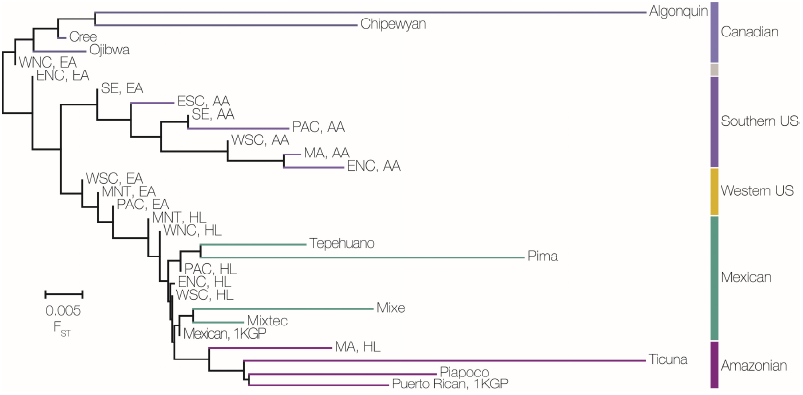
Native American ancestry phylogeny. Phylogenetic relationships are shown for the Native American ancestry-specific components of Native American reference populations (Algonquin, Chipewyan, Cree, Ojibwa, Tepehuano, Pima, Mixe, Mixtec, Ticuna, Piapoco), 1000 Genomes Project reference populations (Mexican and Puerto Rican) and HRS groups. The HRS groups are labeled according to their US census region origins (see Supplementary Figure S3) and genetic ancestry group: African American (AA), European American (EA), and Hispanic/Latino (HL). Broad geographic and genetic groupings are indicated by the bars on the right side. The scale bar corresponds to the pairwise F_ST_ values used to generate the phylogeny.

### Native American ancestry of the *Nuevomexicanos*

The ADMIXTURE results for the Hispanic/Latino group in the Mountain region (MNT) point to the presence of two distinct sub-populations, one of which is clearly Mexican in origin, whereas the second group has a very distinct pattern compared to any other Hispanic/Latino group analyzed here (Figure 4B and Figure 6A). If these two apparent Mountain Hispanic/Latino sub-populations are considered separately, they form distinct phylogenetic groups (Figure 6B). One group clearly falls into the clade with the other Mexican origin populations (see MNT, Mexican), whereas the distinct group is basal to the Mexican clade and intermediate between the Western US and Mexican clades (see MNT, *Nuevomexicano*). The results of the ADMIXTURE and phylogenetic analyses are consistent with historical records indicating the presence of a unique group of Spanish descendants in the American Southwest, known as the ‘Hispanos of New Mexico’ or *Nuevomexicanos*. This population is descended from very early Spanish settlers to the Four Corners region of the US, primarily New Mexico and southern Colorado, and distinct from Mexican-American immigrants who arrived later^27^.

**Figure 6.**
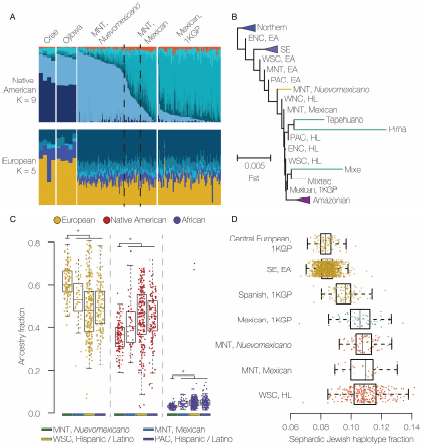
Genetic ancestry of the *Nuevomexicanos*. (A) Native American (*K*=9) and European (*K*=5) ancestry-specific ADMXITURE plots comparing the Mountain census region (MNT) in the middle panel to Native American Cree, Ojibwa (left) and admixed Mexican (right) reference populations. Native American ancestry profiles for the Mountain region can be divided into *Nuevomexicano* (left) and Mexican-American (right) components. (B) Native American ancestry phylogeny (as shown in Figure 5) with the Mountain census region (MNT) broken down into *Nuevomexicano* and Mexican-American components. (C) Distributions of European, Native American, and African ancestry fractions are shown for the Mountain (MNT) *Nuevomexicano*, Mountain (MNT) Mexican, West South Central (WSC) Hispanic/Latino, and Pacific (PAC) Hispanic/Latino groups. The * indicates significant differences in median ancestry fractions between the *Nuevomexicano* and other groups (*P*<0.01 Wilcoxon Rank-Sum test). (D) Distributions of the Sephardic Jewish haplotype copying fractions are shown for European reference populations from the 1000 Genomes Project (Central European and Spanish), HRS European Americans from the Southeast census region (SE, EA), Mountain (MNT) *Nuevomexicano*, Mountain (MNT) Mexican, and West South Central Hispanic/Latino (WSC, HL) groups.

Members of the *Nuevomexicano* population have maintained a distinct cultural identity for centuries, and the ability to isolate individuals from this group based on analysis of their genotypes allowed us to address open questions related to their ancestry. In addition to characterizing their distinct pattern of Native American ancestry, we also compared the levels of Native American admixture between *Nuevomexicanos* and the other nearby Hispanic/Latino groups, which show a more Mexican pattern of Native American ancestry. Consistent with previous results^28^, we show that *Nuevomexicanos* have significantly more European ancestry and less Native American ancestry than other Hispanic/Latino groups from the Western Census regions (Figure 6C). *Nuevomexicanos* also show significantly lower levels of African ancestry compared to the other Hispanic/Latino groups.

*Nuevomexicano* cultural and historical traditions suggest that many of the early Spanish settlers in the region were *Conversos*, or crypto-Jewish individuals, who ostensibly converted to Catholicism in an effort to avoid religious persecution and pogroms, while secretly maintaining Jewish identity and traditions^29^. We interrogated this idea by comparing the extent of Sephardic Jewish admixture found among individuals with the *Nuevomexicano* ancestry pattern compared to other Hispanic/Latino populations. Sephardic Jewish admixture was measured by comparing European haplotypes from Hispanic/Latino individuals to a reference panel including both European and Sephardic Jewish populations. *Nuevomexicanos* show elevated levels of matching to Jewish haplotypes compared to Spanish and other European populations, consistent with substantial *Converso* ancestry among New World Hispanic/Latino populations^30^ (Figure 6d). However, *Nuevomexicanos* do not show a higher level of *Converso* ancestry compared to the other New World Hispanic/Latino populations.

## Discussion

### Native American admixture patterns for distinct US ancestry groups

We were able to delineate three predominant genetic ancestry groups – African American, European American, and Hispanic/Latino – using comparative analysis of whole genome genotypes from >15,000 individuals from across the continental US. Each of these different groups of people experienced distinct historical trajectories in the US, which we found to be manifested as group-specific patterns of Native American ancestry.

Individuals from the African American ancestry group show low (Figure 1B) and relatively invariant (Figure 3A) levels of Native American ancestry across the continental US. The patterns of Native American ancestry seen for the African American group are also more constant among US census regions compared to individuals from the other two ancestry groups (Figure 4A). With respect to the Native American component of their ancestry, African Americans from all US census groups form a single monophyletic clade, for which the Southeast European American (SE, EA) group is basal (Figure 5). Considered together, these results point to a most likely scenario whereby African descendants admixed with local Native American groups in the antebellum South. Early admixture with Native Americans in the South was followed by subsequent dispersal across the US during the Great Migration in the early to mid-twentieth century^31^. The genetic legacy of the Great Migration has previously been explored based on overall patterns of African American genetic diversity^32^. Here, we were able to uncover traces of this same history based solely on the relatively low Native American ancestry component found in the genomes of African Americans.

The European American group shows the lowest levels of Native American ancestry for the three US ancestry groups analyzed here (Figure 1B), consistent with a large and fairly constant influx of European immigrants to the US along with social and legal prohibitions against miscegenation^33^. Compared to African Americans, individuals from the European American ancestry group show more variant levels of Native American ancestry among US census regions (Figure 3B) along with substantially more region-specific patterns of Native American ancestry (Figure 4A). Their region-specific patterns of Native American ancestry are also reflected in the Native American ancestry-based phylogeny, whereby the European American groups are related according to their geographic distribution across the country (Figure 5). These results point to a historical pattern of continuous, albeit infrequent, admixture between local Native American groups and European settlers as they moved westward across the continental US.

As can be expected, the Hispanic/Latino group shows by far the highest (Figure 1B) and most variable (Figure 3C) levels of Native American ancestry across the US. In particular, individuals from the Hispanic/Latino group show highly regional-specific patterns of Native American ancestry (Figure 4B), which are consistent with known demographic trends. For example, analysis of the Native American component of Hispanic/Latino ancestry is sufficient to distinguish Puerto Rican immigrants from the Mid-Atlantic census region from Mexican Americans who predominate in the western census regions. Perhaps most striking, the patterns of Native American ancestry seen for the Mountain census regions were alone sufficient to distinguish descendants of very early Spanish settlers to the region, the group known as Hispanos or *Nuevomexicanos*, from subsequent waves of Hispanic/Latino immigrants who arrived later from Mexico.

The three main US ancestry groups are also distinguished by their patterns of sex-biased ancestry in a way that reflects the unique history of each group (Figure 2). European Americans show very little evidence for sex-biased ancestry, along with very low levels of overall admixture, compared to the African American and Hispanic/Latino groups. The strongest pattern of sex-biased ancestry was seen for the Hispanic/Latino group followed by African Americans. Sex-bias for Hispanic/Latinos is characterized by a strong female-bias for Native American ancestry coupled with European male-biased ancestry. This pattern has been observed in a number of previous studies and is consistent with the history of male-biased migration to the region dating back to the era of the conquistadors^22;34^. The African American group shows female-biased African ancestry and male-biased European ancestry, a pattern which has also been documented previously and tied to the legacy of slavery and racial oppression in the US^35;36^. It has not been previously possible to directly compare the extent of sex-biased admixture among the three largest ancestry groups in the US as we have done here. As such, it is interesting to note that the history of the Spanish colonization in Latin America had a stronger impact on sex-biased ancestry than the legacy of slavery in the US.

### Implications of genetic ancestry for the historical and cultural traditions of *Nuevomexicanos*

Our ability to distinguish *Nuevomexicanos* from the HRS dataset, using their patterns of Native American ancestry, allowed us to address a number of open questions and controversies regarding the history and culture of this interesting population. *Nuevomexicanos* from the American southwest are historically defined as the descendants of early Spanish settlers, those who arrived in the period from 1598 to 1848, as opposed to immigrants from Mexico who arrived the region considerably later. The two distinct patterns of Native American ancestry seen for Hispanic/Latino individuals from the Mountain census region are very much consistent with this historical definition. The *Nuevomexicanos* show a pattern of Native American ancestry that is intermediate to the Canadian and Mesoamerican reference populations analyzed here, whereas the Mexican American individuals from the same region are more closely related to Mesoamerican reference populations. This is consistent with early admixture with local Native American groups in the US southwest, for the *Nuevomexicanos*, versus admixture with Mesoamerican groups in Mexico for the later Mexican immigrants. A more precise characterization of *Nuevomexicanos’* Native American ancestry would require access to genomic data from US Native American reference populations, which are not readily available owing to cultural resistance to genetic testing for ancestry among these groups^37^.

Historically, *Nuevomexicanos* have identified strongly with their European (Spanish) ancestry, while downplaying ancestral ties to Native Americans^38^. This tradition of exclusive European identity is rooted in the colonial era when Spanish descendants in the region were preoccupied with the notion of maintaining so-called pure blood, and the local aristocracy identified as Castilian. Mexicans, on the other hand, have long identified as *Mestizo* with an explicit recognition of their Native American heritage^39^. Our comparative analysis of genetic ancestry for *Nuevomexicanos* and Mexican ancestry groups yielded results that are partly consistent with this historical narrative. On the one hand, *Nuevomexicanos* do have a substantial amount of Native American ancestry, with a median of just under 40% (Figure 6C), which is far more than seen for the African American and European American groups analyzed here, and also more than seen for an number of other Latin American populations in the Caribbean and South America^40^. Nevertheless, the *Nuevomexicanos* have significantly less Native American ancestry, and more European ancestry, than nearby Mexican descendant populations (Figure 6C). Our results are consistent with a recent study that used microsatellite-based ancestry analysis on a much smaller sample of self-identified *Nuevomexicanos*, who were also found to have higher European ancestry and lower Native American ancestry compared to Mexican Americans^28^. Interestingly, we found that the *Nuevomexicanos* also have significantly less African ancestry than Mexican descendant populations, which likely reflects higher levels of early African admixture in Mexico^41^.

Perhaps the most controversial aspect of *Nuevomexicano* history relates to the influence of *Conversos*, or crypto-Jewish individuals, on the culture and traditions of the local community. *Conversos* are Jewish people who converted to Catholicism under intense pressure from religious persecution in Spain, and elsewhere in Europe, and many Spanish *Conversos* immigrated to the New World^42^. Despite their forced conversion to Catholicism, some New World *Conversos* apparently maintained Jewish religious traditions over the centuries since their immigration from Spain. For example, the persistence of rituals and symbols related to Jewish traditions in New Mexico has been taken as evidence for an influential presence of *Conversos* among the *Nuevomexicanos*, a position championed by the historian Stanley Hordes^29^. On the other hand, the folklorist Judith Neulander and others have been fiercely critical of this narrative based on what they perceive to be misunderstandings of the origins of many of the cultural traditions tied to Jewish rituals and even deliberate misrepresentations of facts^43^. Neulander’s interpretation relates the notion of *Converso* identity among *Nuevomexicanos* back to the colonial assertions of pure Spanish ancestry given that the Sephardim are Spanish and would presumably be loath to marry outside of their religious group^44^.

We evaluated the extent of Sephardic Jewish ancestry among *Nuevomexicanos*, via comparative analysis of their European haplotypes to both European and Sephardic Jewish reference populations, in attempt to assess the genetic evidence in support of the *Converso* narrative. While we did find more Sephardic Jewish ancestry among *Nuevomexicanos* compared to Spaniards or other Europeans, they did not show any more Sephardic Jewish ancestry than Mexican descendants from nearby regions (Figure 6D). Our results are consistent with a recent study that used haplotype-based ancestry methods to uncover widespread *Converso* ancestry in Latin American populations^30^. Taken together, we interpret these results to indicate that, while *Nuevomexicanos* do in fact have a demonstrable amount of Jewish ancestry, they are no more, or less, Jewish than other New World Latin American populations. Of course, we cannot weigh in on the strength of evidence for or against the persistence of Jewish cultural traditions among *Nuevomexicanos* based on our genetic evidence alone. Nevertheless, there does not seem to be anything particularly unique, at least from the genetic perspective, with respect to the extent of Sephardic Jewish heritage among *Nuevomexicanos*.

## Conclusion

Much of the genetic legacy of the original inhabitants of the area that is now the continental US can be found in the genomes of the descendants of European and African immigrants to the region. In this study, we analyzed signals of cryptic Native American genetic ancestry that can be gleaned from comparative analysis of genomes from three distinct US ancestry groups: African American, European American, and Hispanic/Latino. Our study was enabled by the use of haplotype-based methods for genetic ancestry inference and leveraged a large dataset of whole genome genotypes. This approach allowed for detailed analysis of Native American ancestry patterns even when the per-genome levels of Native American ancestry were quite low, *i.e.* cryptic genetic ancestry. Each of the three genetic ancestry groups analyzed here shows distinct profiles of Native American ancestry, which reflect population-specific historical patterns of migration and settlement across the US. Analysis of the Native American ancestry component for members of these groups allowed for the delineation of region-specific subpopulations, such as the *Nuevomexicanos* from the American southwest, and facilitated the interrogation of specific historical scenarios.

